# Compensatory adaptation does not fully alleviate the costs associated with rifampicin resistance and occurs predominantly through off-target mutations

**DOI:** 10.1101/2023.01.13.524028

**Authors:** Yasmin Cohen, Gydeon A Rozen, Pleuni S Pennings, Sophia Katz, Ruth Hershberg

## Abstract

The manner in which costs associated with antibiotic resistance mutations are alleviated through the acquisition of compensatory mutations has bearing on the dynamics of resistance accumulation and spread as well as on our more general understanding of the persistence of adaptive genotypes. Here, we combine evolutionary experiments, fitness analyses, and whole genome sequencing to examine the way compensation to alleviate costs associated with rifampicin resistance is achieved, both when resistance is fixed and when initial majorities of resistant cells are grown alongside susceptible cells. We found compensatory mutations to occur across all examined populations. However, compensation never fully alleviates the initial costs of resistance. In contrast to prevalent assumptions, compensatory mutations mostly occurred outside of the RNA polymerase core enzyme, which serves as the target of rifampicin. The target size for compensatory mutations appears to be high, as great variation exists in compensatory mutations, even within a single population. At the same time, the pathways of compensation are quite convergent, meaning that, across independent populations, we often observe compensatory mutations in the same loci and sometimes even observe the precise same compensatory mutations across populations.

## Introduction

The development of antibiotic resistance is among the most well-studied types of adaptation, largely due to its great medical importance (e.g. (1-7)). By studying the dynamics of adaptation to antibiotics, one can forward our general understanding of how adaptation works while at the same time gaining knowledge needed to develop the best strategies for combating the accumulation and spread of resistance.

Bacteria can achieve resistance to antibiotics through point mutations within the gene targets of the antibiotic or through the acquisition of novel genes through horizontal gene transfer. Antibiotics are designed to target important bacterial functions such as replication, transcription, and translation. Such important functions are generally regulated by the most conserved housekeeping genes, which thus often serve as antibiotic targets (8). Point mutations that lead to antibiotic resistance often do so by altering the antibiotic binding site on their gene targets, so the antibiotic can no longer induce their harmful effect on the bacteria. Because antibiotics so often target central housekeeping regulatory genes, resistance mutations often fall within these highly constrained genes, leading to substantial costs to bacterial fitness (reviewed in (9, 10)). Such costs manifest most strongly when these bacteria are no longer exposed to the antibiotics to which these mutations confer resistance. It was therefore hoped that reversion mutations leading to loss of resistance would be favored by selection once treatment with antibiotics stops (9). Instead, studies have shown that bacteria can rapidly alleviate the costs of antibiotic resistance mutations through the acquisition of compensatory mutations while maintaining the resistance phenotype (e.g., (9, 11-13)). Such compensatory mutations tend to occur more frequently than reversion mutations because, for each resistance mutation, there is only one potential reversion mutation but several potential compensatory mutations. For this reason, several studies have shown that in lab populations in which resistance is initially fixed, resistance will tend to be maintained over the time frames examined (usually on the range of hundreds of generations) rather than lost, even once populations are no longer treated with antibiotics (9, 11, 12). A more recent theoretical study (14), which was motivated and supported by experimental results on compensation in the context of adaptation to prolonged resource exhaustion (15), suggests that under normal mutation rates, resistance will indeed tend to persist. However, unless compensatory mutations are able to alleviate the costs of resistance fully, increases in mutation rate will tend to lead to much more frequent reversion (14).

The antibiotic rifampicin targets the bacterial RNA polymerase core enzyme gene *rpoB* and resistance to rifampicin is thus achieved through mutations to *rpoB* (16). While rifampicin is not important in the clinical treatment of *Escherichia coli*, it is important in the treatment of other pathogens and, most importantly, in the treatment of tuberculosis. Studies of compensatory evolution have been carried out quite broadly using rifampicin resistance as a model, utilizing both lab experiments in model bacteria such as *E. coli* and *Salmonella*, and also examining the patterns of compensatory mutation accumulation in *Mycobacterium tuberculosis* (e.g. (12, 17-21). These studies often tend to focus only on the genes of the RNA polymerase core enzyme as potential targets for compensation.

Antibiotic resistance is often assumed to be fixed within antibiotic-resistant populations of bacteria, as the selective pressure exerted by the presence of antibiotics is thought of as lethal to non-resistant bacteria. For this reason and for simplicity, most of the experiments aimed at understanding the dynamics of post-resistance evolution are carried out on populations in which resistance is, at least initially, fixed. However, due to antibiotic persistence and incomplete penetrance of antibiotics, non-resistant cells may be able to survive antibiotic treatment within the bodies of hosts (22, 23). As a result, antibiotic resistance may not become fixed, even within populations of bacteria that become largely resistant in response to antibiotic treatment. In populations in which an adaptive yet costly genotype is not entirely fixed, fluctuations in genotype frequencies can occur, once conditions no longer favor the adaptive genotype, leading to rapid reductions in its frequency (15). The extent to which this may lead to reductions in antibiotic resistance frequencies will likely depend on the sizes of the minorities of the susceptible cells, on the costs of resistance mutations, and on the rates with which compensatory evolution can occur to alleviate these costs. On the flip side of this, the dynamics of compensatory evolution may vary between populations in which resistance is entirely fixed and populations in which resistant cells need to compete against co-existing susceptible cells.

Here, we carried out a set of evolutionary experiments using the model bacterium *Escherichia coli* to examine the manner in which both populations fixed for rifampicin resistance and populations in which resistant clones are initially mixed with small minorities of susceptible cells evolve to compensate for costs associated with antibiotic resistance. We follow up our evolutionary experiments with whole genome sequencing of clones from each evolved population and with competition experiments to estimate the effects of compensation on fitness. We show that compensation occurs in both resistance-fixed and resistance-mixed populations, although costs are never fully alleviated. We further show that compensation occurs in a largely convergent manner across populations, through similar mutations to similar loci, and that, contrary to widespread assumptions, compensation only rarely occurs through mutations to the rifampicin target gene complex and most often occurs off-target.

## Results

### Rapid fluctuations in resistance frequencies are observed in populations in which large majorities of antibiotic-resistant cells are grown along minorities of susceptible cells

In this study, we focused on a single rifampicin-resistant mutant carrying a RpoB S512Y mutation. This mutant was fully sequenced to verify that it is isogenic to its ancestral *E. coli* K12 MG1655 genotype, except for the one resistance mutation (3). The S5212Y mutant was selected because, in contrast to some other similarly isogenic rifampicin-resistant mutants, it did not carry a striking phenotypical cost when it was independently grown in the absence of rifampicin (**Figure 1A**). We could thus conclude that the fitness cost of this mutation was not extreme relative to that of other rifampicin-resistance mutations.

**Figure 1.**
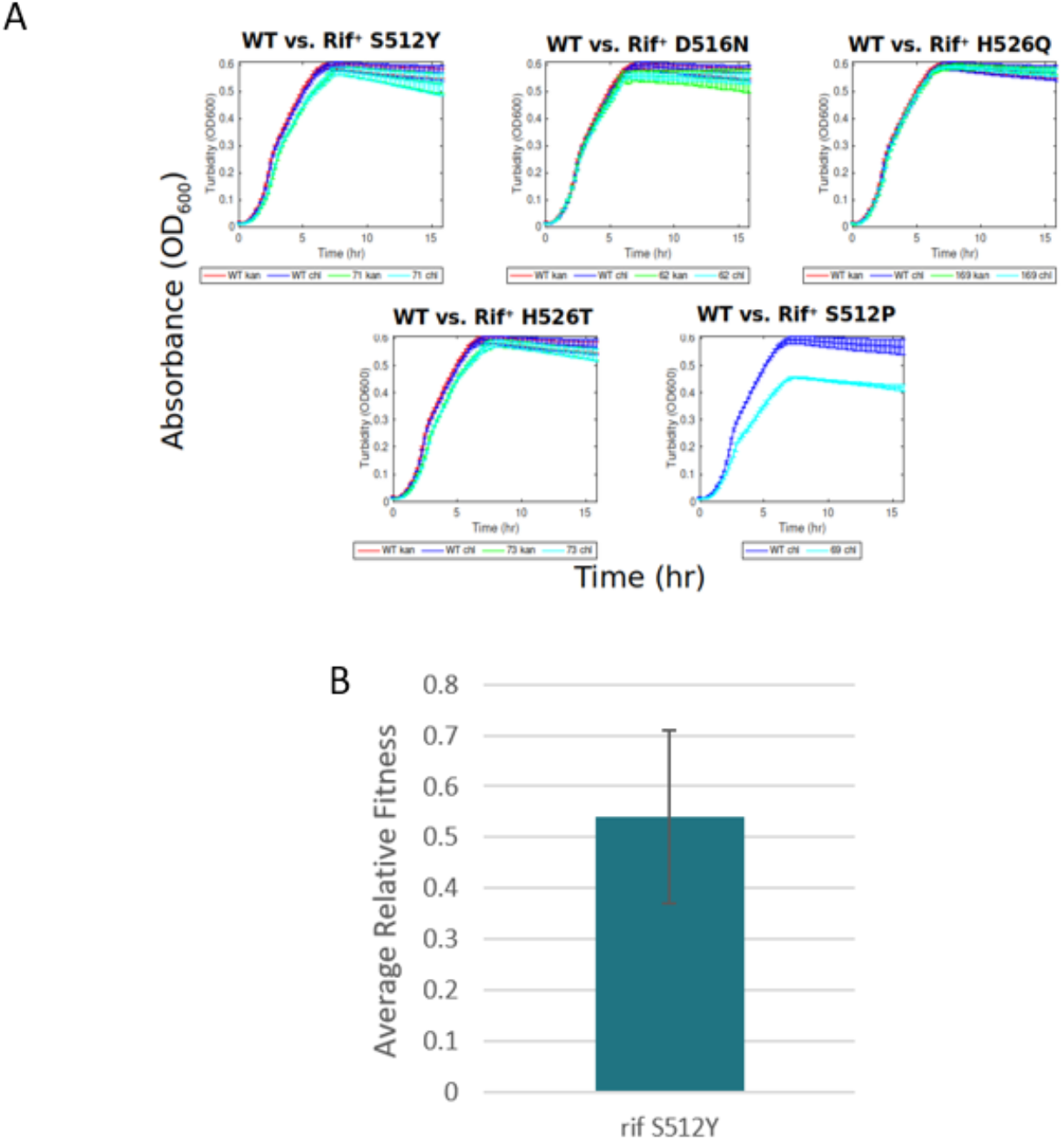
Fitness measurements for the ancestral rifampicin resistant RpoB D512Y mutant used in this study. **(A)** the RpoB D512Y ancestral genotype does not show a pronounced effect on growth rates when grown by itself in rich LB media. Depicted are growth curves observed through measurements of OD in a plate reader for five different rifampicin-resistant genotypes that were fully sequenced and found to carry only the single resistance mutation. Pale blue lines represent independent measurements for each resistant strain, while purple lines represent the same for the ancestral susceptible *E. coli* K12 MG1655 genotype (grown in the same plate reader experiments in different wells). (**B**) The RpoB S512Y mutant does suffer a substantial reduction in fitness relative to its ancestral susceptible genotype when competed together (see also **Table 1 and Table S1**).

We generated strains marked with either Kanamycin (Kan) or Chloramphenicol (Chl) resistance cassettes for both the WT and the RpoB S512Y mutant (3). Competition experiments were carried out using these strains to characterize this mutant’s relative fitness against its ancestral susceptible (WT) strain. Remarkably, despite having no observable effect on independent growth, the RpoB S512Y mutant suffers a substantial cost to its relative fitness (w) (w = 0.54, **Figure 1B, Table 1** and **Table S1**, w is significantly lower than 1 (*P* < 0.001, according to a one-tailed paired Mann Whitney test)).

**Table 1.** Results of competition experiments for evolved population samples

| population/clone | Fitness test 1<br>(relative WT<br>ancestor)<sup>a</sup> | P-value for<br>test 1<sup>b</sup> | Fitness test 2 (relative<br>Ancestral Rifampicin<br>resistant RpoB S5212Y<br>mutant)<sup>a</sup> | P-value for<br>test 2<sup>c</sup> |
| --- | --- | --- | --- | --- |
| Ancestral<br>Rifampicin<br>resistant RpoB<br>S5212Y mutant | **0.54** +/- 0.17<br>(n=11) | N/A | N/A | N/A |
| Evolved 90_kan_1<br>population | **0.73** +/- 0.06<br>(n=10) | 0.004 | **1.14** +/- 0.19<br>(n=12) | 0.026 |
| Evolved 99_kan_1<br>population | **0.63** +/- 0.09<br>(n=11) | 0.179 | **1.11** +/- 0.14<br>(n=15) | 0.009 |
| Evolved 100_chl_1<br>population | **0.83** +/- 0.12<br>(n=12) | <0.001 | **1.15** +/- 0.12<br>(n=10) | <0.001 |

<sup>a</sup>For each type of competition carried out, we present the mean value of the relative fitness obtained (see Materials and methods) in bold, +/- the value's standard deviation around the mean, and (n=the number of competition experiments carried out).
<sup>b</sup>Mann-Whitney non-paired one-tailed tests were carried out to distinguish: H0: Relative fitness against the wild type for the evolved rifampicin resistant population <= Relative fitness against the wild type for the ancestral rifampicin resistant RpoB S5212Y mutant; and H1: Relative fitness against the wild type for the evolved rifampicin resistant population > Relative fitness against the wild type for the ancestral rifampicin resistant RpoB S5212Y mutant. The resulting P-values are provided.
<sup>c</sup>Mann-whitney paired one-tailed tests were carried out to distinguish: H0: Relative fitness of evolved rifampicin resistant population against the ancestral rifampicin resistant RpoB S5212Y mutant <=1; and Relative fitness of evolved rifampicin resistant population against the ancestral rifampicin resistant RpoB S5212Y mutant >1.

Using the Kan- and Chl-marked ancestral susceptible (WT) and RpoB S512Y rifampicin resistant strains, we initiated four sets of 16 cycle 1:100 serial dilution evolutionary experiments (see **Materials and methods**) starting with either: (1) 100% WT susceptible cells; (2) 100% RpoB S512Y rifampicin resistant cells; (3) 99% rifampicin-resistant and 1% susceptible cells; or (4) 90% rifampicin resistant cells and 10% susceptible cells. Serial dilution experiments were carried out in Luria broth (LB), containing no antibiotics. Each day bacteria were diluted 1:100 into fresh LB. Each of the four sets of experiments included four independent populations (for a total of 16 populations). For each set of four experiments, in two, the resistant cells were marked with Kan and the susceptible ones with Chl, and in the other two, the resistant cells were marked with Chl and the susceptible cells with Kan. Following 16 days of serial dilution (∼100 generations), we measured what fraction of the endpoint cells carry resistance to rifampicin. Entirely susceptible populations evolved resistance to rifampicin at an average frequency of ∼10^−8^ (**Figure 2**). Populations starting off with 100% resistant cells maintained ∼100% resistance and did not seem to undergo substantial reversion. In contrast, populations initiated with minorities of susceptible cells underwent pronounced reductions in rifampicin-resistant cell frequencies, which reduced from 0.99 or 0.9 to between 7*10^−8^ and 5*10^−3^ (**Figure 2**). By also quantifying the frequency with which cells carry resistance to Kan or Chl, we could show that, within mixed populations, the frequency of rifampicin resistance always matched very well the frequency of the marker that marked the resistant cells that were initially inoculated into each population (**Figure 2**). This demonstrates that the reduction in resistance frequencies occurred largely through fluctuations in genotype frequencies rather than due to reversion mutations. Despite initially mixed populations undergoing very rapid reductions in resistance frequencies, resistance was never entirely lost from any of the studied populations.

**Figure 2.**
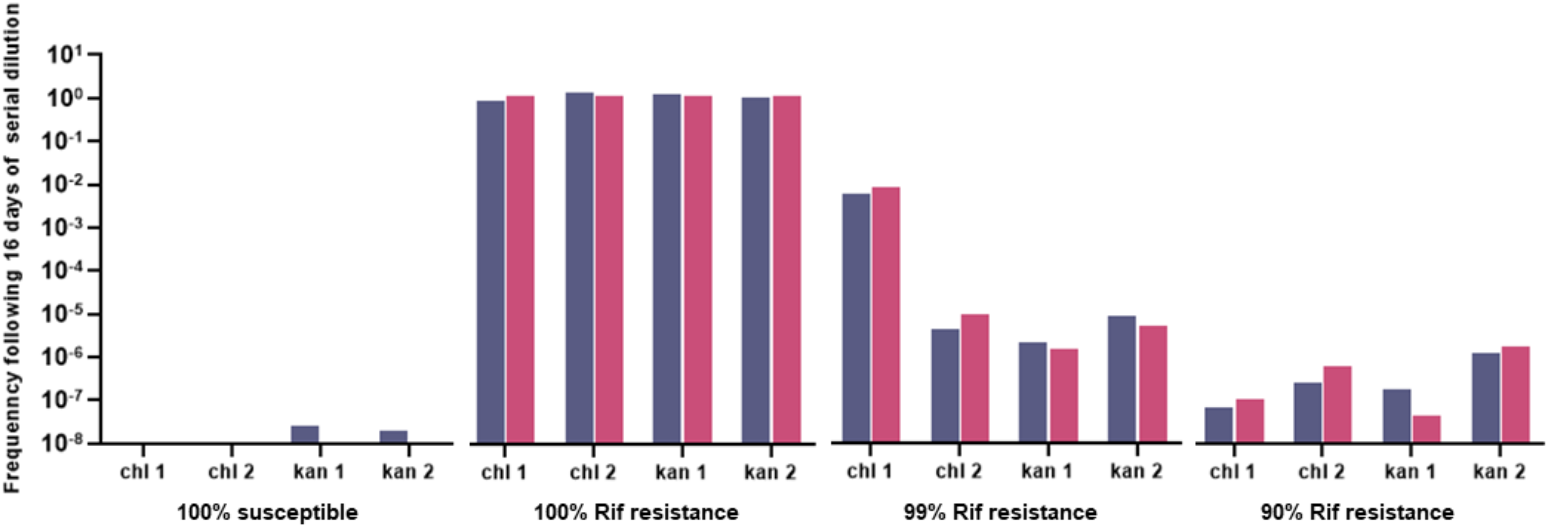
Ultimate frequencies of rifampicin-resistant cells within 16 populations of *E. coli*, following 16 daily cycles of serial dilution (∼100 generations of growth) within LB containing no antibiotics. Populations that were initially fixed for resistance did not suffer reductions in resistance frequencies due to reversion. The initial presence of 1% to 10% minorities of susceptible cells within populations of largely rifampicin-resistant bacteria lead to rapid reductions in resistance frequency in the absence of rifampicin exposure. Depicted are for each one of the 16 serial dilution experiments the frequency of rifampicin-resistant cells (gray) and the frequency of cells carrying the marker (kan or chl) that originally marked the resistant members of the population (pink) following 16 days of serial dilution. For the eight populations that originally contained both resistant and susceptible cells, the eventual frequency of rifampicin resistance matches very well the eventual frequency of the selective marker (kan or chl) that originally marked the rifampicin-resistant cells. This indicates that the reduction in the frequency of rifampicin resistance was mostly due to fluctuations in genotype frequencies rather than due to reversion mutations.

### Compensatory evolution occurs within evolved populations, yet compensation never fully alleviates the initial costs of resistance

Next, we selected at random one of the serially diluted populations that initially had 100% resistant cells (from here on referred to as 100_chl_1), one that initially had 99% resistant cells (99_kan_1), and one that initially had 90% resistant cells (90_kan_1). For each such population, we aimed to examine whether resistant cells remaining within that population, following 16 days of serial dilution, compensated for the initial costs of resistance. To do so, we carried out two types of competition experiments: (1) against the wildtype ancestral susceptible strain – if the relative fitness of an evolved rifampicin-resistant population or clone against the wildtype is significantly higher than the relative fitness of the ancestral rifampicin-resistant genotype against the same wildtype, we can deduce that some compensation has occurred. (2) Competitions against the ancestral resistant strain – here, we can say compensation has occurred if the relative fitness of the evolved rifampicin-resistant population or clone against its ancestral resistant genotype is significantly higher than 1.

All three populations showed significantly enhanced fitness relative to their ancestral rifampicin-resistant genotype (**Table 1, Table S2**). Two of the three evolved rifampicin-resistant populations also statistically improved their relative fitness vs. the ancestral wildtype significantly, beyond what we saw in the original competition between their ancestral resistant strain and the wildtype (**Table 1, Table S1**). For the third population (99_kan_1), we did see a higher mean fitness, but this increase was not statistically significant. At the same time, none of the populations fully alleviated the costs of resistance, as all of them were still significantly less fit than the wildtype (w against wildtype ancestor significantly lower than 1, *P* < 0.001, according to paired single-tailed Mann-Whitney test).

### Convergent yet diverse compensatory mutations occur mostly off-target

In order to understand how compensation was achieved, we fully sequenced ∼10 rifampicin-resistant clones out of each of the 12 evolved populations that were initiated with either 100%, 99%, or 90% resistant cells (for a total of 118 fully sequenced clones). The coverage obtained for each clone is summarized in **Table S3**. A full list of identified mutations is provided in **Table S4**.

As expected, all resistant clones sequenced carried the original RpoB S512Y mutation. In addition, for 107 (90.7%) of the sequenced clones, we could identify at least one secondary, putatively compensatory mutation that arose on the background of the original RpoB resistance mutation (**Figure 3, Table S4**). Surprisingly, putatively compensatory mutations very rarely occur within RpoB. Of the 107 clones carrying putatively compensatory mutations, the mutation fell within RpoB in only 12 clones (11.2%). Clones with RpoB mutations were seen in only two of the 12 populations, and in both these populations, the exact same mutation occurred (RpoB P1044L). No other mutations were seen in genes encoding the RNA polymerase core enzyme.

**Figure 3.**
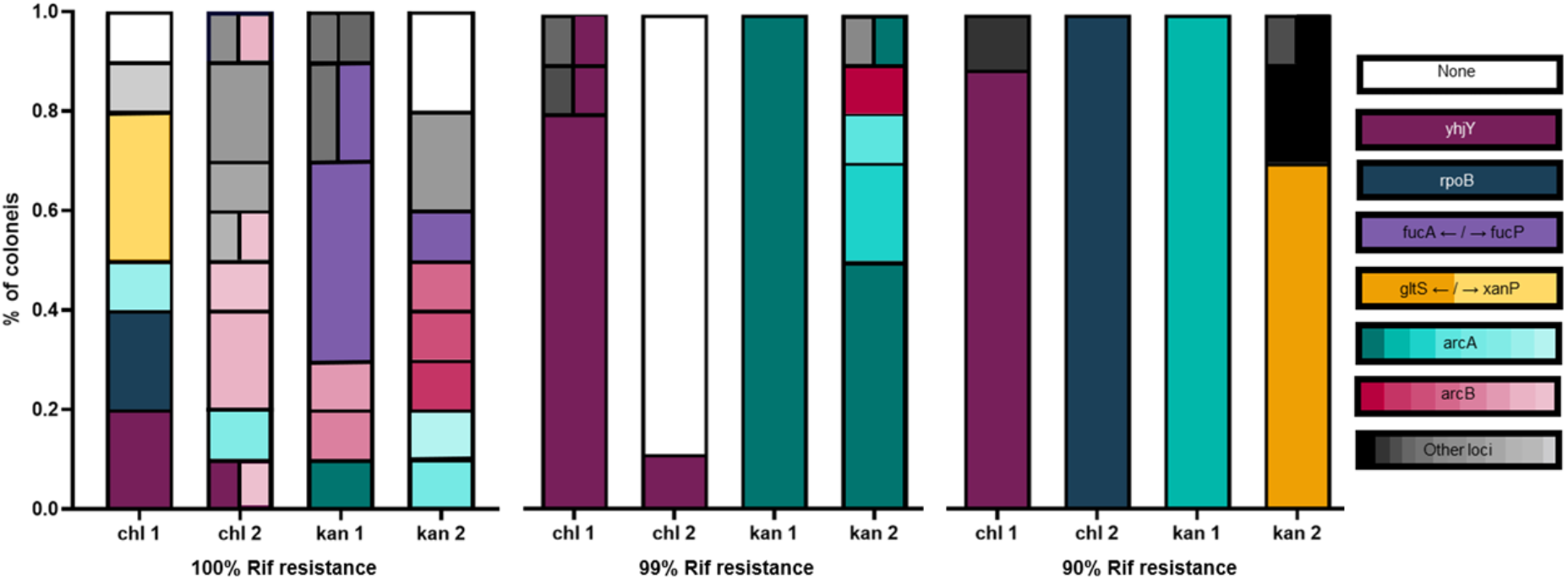
Summary of putative compensatory mutations identified by sequencing the entire genomes of ∼10 clones from each of the 12 populations that initially contained resistant clones following 16 days of serial dilution (∼100 generations of growth). All sequenced rifampicin-resistant clones maintained within them the original RpoB D512Y rifampicin resistance mutation. Loci in which mutations were observed in multiple populations were designated their own color palate, with each specific mutation being given its own distinct shade within that color palate. White represents clones in which no compensatory mutations were observed. A grayscale color palate designates loci in which only a single mutation was found within only a single population. *gltS* **←** / **→** *xanP* refers to the intergenic region between the *gltS* and *xanP* genes, and *fucA* **←** / **→** *fucP* refers to the intergenic region between the *fucA* and *fucP* genes.

Other genes in which putatively compensatory mutations occurred across multiple populations include *arcA* (31.8% of putatively compensated clones, across seven of the 12 populations), *arcB* (11.2% of putatively compensated clones, across four of the 12 populations), *yhjY* (20.6% of putatively compensated clones, across five of the 12 populations), the intergenic region between the *fucA* and *fucP* genes (7.5% of putatively compensated clones, across two of the 12 populations), and the intergenic region between the *gltS* and *xanP* genes (9.3% of putatively compensated clones, across two of the 12 populations). In the cases of *yhjY*, and the intergenic region between *fucA* and *fucP*, precisely the same mutations were found across all independently evolved populations in which mutations occurred within these loci. For *arcA*, seven different mutations were found within the gene or its promoter region in the five populations in which mutations were seen within this locus. Nevertheless, one specific mutation within *arcA* (F79L) was seen across three independently evolved populations. For *arcB*, eight different mutations were observed in the four populations that contained clones carrying mutations within *arc*B.

While most clones acquired only a single putatively compensatory mutation, ten of the sequenced clones acquired two different putatively compensatory mutations (**Figure 3**). In most such instances, one or both mutations did not fall within genes that were mutated in a convergent manner across multiple independently evolving populations. It is thus difficult to predict whether both mutations in such instances are indeed compensatory, as it is quite possible that one or both of them are simply neutral passenger mutations. However, we did see one clone in which two mutations were acquired in loci that were both mutated convergently across additional populations (*arcB* and *yhjY*). Such convergence suggests that mutations within these loci occur because they provide a benefit, and it is thus likely that this clone has acquired two different compensatory mutations in the relatively short timeframe of our evolutionary experiment.

Many populations ended up with a diversity of putatively compensatory mutations, with different clones from the same population often carrying different mutations. A clear pattern emerged by which evolved populations initiated from 100% rifampicin-resistant cells ended up with a much higher variation of putatively compensated genotypes compared to those populations that were initiated with 99% or 90% of resistant cells (**Figure 3**). For populations initiated with 100% resistant cells, the mean number of different putatively compensatory mutations observed following 16 days of serial dilution was 6.5 +/- 1.3. At the same time, this number was 2.5 +/- 1.9 for populations initiated with 99% resistant cells and 1.75 +/- 1 for populations initiated with 90% resistant cells.

The great convergence in the loci mutated across populations suggests that many of the mutations observed benefit fitness and are thus compensatory. However, we sought to test this more directly. Competition experiments were carried out to examine whether the putatively compensatory mutations we identified are indeed compensatory (i.e., whether they improve the relative fitness of the clones carrying them). For these experiments, we again focused on the three randomly selected populations analyzed above. We carried out competition experiments between each of the putatively compensated genotypes that evolved within these populations and the ancestral susceptible (WT) and ancestral rifampicin-resistant strains (**Table 2**). All but one of the clones examined demonstrated a fitness improvement in at least one of the two types of competition experiments we carried out. Six of the seven putatively compensated clones tested significantly improved their fitness relative to the wildtype ancestral susceptible genotype, compared to the ancestral resistant genotype (**Table 2** and **Table S1**), while three of the seven putative compensatory mutations showed increased fitness in direct competition with the ancestral resistant strain (**Table 2**, and **Table S2**). The only putative compensatory genotype that did not show any significant fitness improvement in either of the two tests used was YhjY S21S. However, it is important to note that this mutation is extremely convergent, occurring in five independently evolving populations (**Figure 3**). It thus seems that it is quite likely that this mutation does indeed improve fitness, despite our inability to measure this improvement directly via competition experiments (see **Discussion** below).

**Table 2.** Results of competition experiments for evolved individual clones

| Clone | Fitness test 1<br>(relative WT<br>ancestor)<sup>a</sup> | P-value for<br>test 1<sup>b</sup> | Fitness test 2 (relative<br>Ancestral Rifampicin<br>resistant RpoB S5212Y<br>mutant)<sup>a</sup> | P-value for<br>test 2<sup>c</sup> |
| --- | --- | --- | --- | --- |
| Ancestral Rifampicin<br>resistant RpoB S5212Y<br>mutant | **0.54** +/- 0.17<br>(n=11) | N/A | N/A | N/A |
| 100_chl1_YhjY_S21S | **0.57** +/- 0.1<br>(n=10) | 0.39 | **1** +/- 0.07<br>(n=10) | 0.754 |
| 100_chl1_RpoB_P1044L | **0.72** +/- 0.06<br>(n=10) | 0.004 | **1.04** +/- 0.06<br>(n=10) | 0.07 |
| 100_chl1_ArcA_N116T | **0.77** +/- 0.07<br>(n=10) | <0.001 | **1.03** +/- 0.09<br>(n=10) | 0.4 |
| 100_chl1_YacC/CueO<br>intergenic region | **0.68** +/- 0.07<br>(n=12) | 0.02 | **1.15** +/- 0.12<br>(n=10) | 0.002 |
| 100_chl1_GltS/XanP<br>intergenic region | **0.72** +/- 0.14<br>(n=10) | 0.013 | **0.9** +/- 0.15<br>(n=12) | 0.96 |
| 90_Kan1_ArcA_G96A<sup>d</sup> | **0.73** +/- 0.06<br>(n=10) | 0.004 | **1.14** +/- 0.19<br>(n=12) | 0.026 |
| 99_Kan1_ArcA_F79L<sup>d</sup> | **0.63** +/- 0.09<br>(n=11) | 0.179 | **1.11** +/- 0.14<br>(n=15) | 0.009 |

<sup>a</sup>For each type of competition carried out, we present the mean value of the relative fitness obtained (see Materials and methods) in bold, +/- the value's standard deviation around the mean, and (n=the number of competition experiments carried out).
<sup>b</sup>Mann-Whitney non-paired one-tailed tests were carried out to distinguish: H0: Relative fitness against the wild type for the evolved rifampicin resistant population <= Relative fitness against the wild type for the ancestral Rifampicin resistant RpoB S5212Y mutant; and H1: Relative fitness against the wild type for the evolved rifampicin resistant population > Relative fitness against the wild type for the ancestral rifampicin resistant RpoB S5212Y mutant. The resulting P-values are provided.
<sup>c</sup>Mann-whitney paired one-tailed tests were carried out to distinguish: H0: Relative fitness of evolved rifampicin resistant population against the ancestral rifampicin resistant RpoB S5212Y mutant <=1; and Relative fitness of evolved rifampicin resistant population against the ancestral rifampicin resistant RpoB S5212Y mutant >1.
<sup>d</sup>Because the rifampicin-resistant sub-population in population 90\_Kan1 and population 99\_Kan1 were fixed for a single rifampicin-resistant mutant, the results for the clone from that population are exactly the same as the results for the population.

In competitions against the susceptible ancestor, all strains still displayed significantly reduced relative fitness, demonstrating that none of the putatively compensatory mutations fully alleviated the costs associated with resistance (*P* << 0.001, **Table 2, Table S1**).

## Discussion

Here, we combine an evolution experiment with competition experiments and whole genome sequencing of individual compensated clones to examine the dynamics of compensatory evolution in populations initially fixed for antibiotic resistance and populations in which initial majorities of resistant cells co-exist with minorities of susceptible cells. We found that when resistance was initially fixed, it remained fixed throughout the ∼100-generation evolutionary experiments we performed. When resistance was not fixed, the frequency of resistant cells sharply decreased, yet nevertheless, resistant cells remained within the population at low frequencies. Both fixed-resistance populations and mixed-resistance populations compensated for the initial costs of resistance by acquiring compensatory mutations. However, such compensatory evolution never fully alleviated the initial costs of resistance, and resistant cells were left with substantial costs to their fitness relative to their ancestral susceptible genotype.

A previous modeling study has shown that when compensatory mutations cannot fully alleviate the costs of resistance, the persistence of resistance should depend on mutation rates (14). In populations where mutators elevate mutation rates, reversions will tend to occur quite rapidly, leading to loss of resistance (14, 15). However, in non-mutator populations, of the type we studied here, we do not expect to see a rapid loss of resistance due to reversion, even when compensatory mutations do not fully alleviate costs. Our current results thus fit with and support these prior modeling-based results. It is possible that the fact that our populations never fully alleviate costs associated with resistance would ultimately, given sufficient time, lead to loss of resistance. However, this is not necessarily the case, as the accumulation of compensatory and other mutations with time may close down the option of reversion as a way to fully alleviate costs due to the effects the compensatory and other mutations may themselves have on fitness. Further studies will be needed to examine the long-term consequences of incomplete compensation on the persistence of antibiotic resistance.

While the initial presence of minorities of susceptible cells alongside the majority of antibiotic-resistant cells within our populations led to rapid reductions in resistance frequencies due to fluctuations in genotype frequencies, they did not lead to a complete loss of resistance. To examine whether this is surprising, given the measured fitness costs suffered by both the ancestral RpoB S512Y rifampicin-resistant genotype and its compensated evolved genotypes, we carried out simulations (Materials and Methods and Supplementary text S1). The results of these simulations are summarized in **Figure S1** and show that given the costs we measured for two of the mixed-resistance populations (**Table 1**), we might naively expect to, at least in some instances, observe a complete loss of resistance. The fact that we never observe such a loss of resistance may thus indicate that frequency-dependent selection or another form of balancing selection is at play. It would be interesting to further investigate this possibility in future studies.

We utilized two types of competition experiments to estimate whether different populations/clones compensated for the initial costs of resistance. In the first type of experiment, we competed the evolved rifampicin-resistant population/clone of interest against the ancestral susceptible genotype. We then examined whether the obtained relative fitness improved in comparison to that seen for the ancestral rifampicin-resistant genotype. In the second type of experiment, we competed the evolved rifampicin-resistant population/clone directly against its ancestral rifampicin-resistant genotype. We could then examine whether this resulted in a relative fitness significantly higher than 1. We consider any type of improvement observed, be it in the first or second test, as evidence for compensatory evolution. It is, however, worth noting that the two tests often do not agree, with only one of them demonstrating a significant improvement in the fitness of the evolved rifampicin-resistant populations/clones. Additionally, we were unable to see a significant increase in fitness using either test for a clone carrying the YhjY S21S putative compensatory mutation, despite this mutation occurring in an extremely convergent manner across five independently evolving populations. Such apparent discrepancies may stem from the noisiness of the measurements themselves but likely often stem from the complexity of relative fitness itself (24). Relative fitness is likely strongly dependent on context and is the result of a complex interplay of factors. As a result, it is quite possible to be able to detect an increase in fitness when competing genotypes against some backgrounds but not others.

Previous studies regarding the manner in which bacteria compensate for the deleterious effects of rifampicin resistance mutations on bacterial fitness have focused mostly on on-target compensatory mutations occurring within the RNA polymerase core enzyme (RNAPC) genes, *rpoA, rpoB, and rpoC* (e.g., (12, 17-21, 25, 26). This stems from an assumption that the simplest way to “fix” a deleterious effect of a mutation within a specific gene or complex is to make additional alterations to that gene or complex and from technical limitations that make it much easier to identify compensatory mutations within a defined gene or complex. Here we find that the vast majority of compensatory mutations that occur within our populations occur outside of the target gene complex of the antibiotic rifampicin. There is no reason to assume that off-target compensatory mutations do not often contribute to compensation for the costs of rifampicin resistance in additional bacteria, including the most clinically relevant *M. tuberculosis*. Indeed, despite receiving less attention, some putative off-target compensatory mutations were already identified in *M. tuberculosis* (27). It is also quite likely that off-target compensatory mutations play a substantial role in the case of resistance to additional antibiotics beyond rifampicin and, more generally, in alleviating the costs of non-antibiotic-related adaptations. Fully understanding the manner in which adaptive costs are compensated will, therefore, likely require a broader focus, not only on the gene or complex in which adaptations occur but on the entire genome.

Our results also shed light on the great diversity of compensatory mutations that develop, over relatively short periods of time, even within a single population. Such diversity was particularly high for populations in which resistance was initially and remained fixed. That carrying a costly resistance mutation leads to the generation of a diversity of compensatory mutations within a single population exemplifies how adaptation to a specific stressor or condition can push forward the generation of variation within a population. Bacteria carrying a costly adaptation need to further adapt to compensate for its initial costs. Various individuals within a single population can acquire different compensatory mutations, leading to increases in variation within that population. These compensatory mutations may also affect the trajectory of which future mutations would be adaptive or deleterious, which may, in turn, lead to the generation of even greater variation with time, in what would constitute an example of historical contingency (28).

The high diversity observed in the identity of compensatory mutations within our populations strongly suggests that the target size for such compensation is fairly large. However, we did find high levels of convergence, with similar loci being mutated in independently evolving populations and sometimes even precisely the same compensatory mutations being observed in several populations. Thus, it is quite likely that while the target size for compensation is quite large and extends far beyond the target genes of antibiotics, there are specific pathways to compensation that are often “hit.” We can therefore hope that despite the diversity of possible compensatory mutations, which often occur outside of an antibiotic’s target gene, it should prove possible to ultimately identify the majority of pathways by which bacteria evolve to compensate for the costs of antibiotic resistance.

## Materials and Methods

### Evolutionary experiments

All experiments were started using WT *E. coli* K12 MG1655 and its derived RpoB S512Y rifampicin-resistant bacteria (isolated in our previous study (3)). Both the WT and rifampicin-resistant strains were previously marked with either Kanamycin (Kan) or Chloramphenicol (Chl) resistant cassettes, generating the four types of strains used: (WT marked with Kan, WT marked with Chl, RpoB S512Y rifampicin-resistant marked with Kan, and RpoB S512Y rifampicin-resistant marked with Chl).

Each type of strain was extracted from its glycerol stock, streaked on Luria broth (LB) agar plates, and left to grow upside down overnight at 37°C in an incubator. Following overnight growth, a single colony was sampled from each plate and inoculated into 4 ml of fresh LB in 14 ml tubes and transferred to a shaking incubator (225 rpm) for 2 hours at 37°C until an approximate OD of 0.2 was reached. All 0.2 OD_600_ cultures were diluted five-fold (1:5, v/v), and the non-diluted (1:1) and diluted (1:5) samples were further appropriately diluted into LB broth to achieve a final one-hundred-fold (1:100, v/v) dilution in a final volume of 4 ml LB in 14 ml test tubes, thereby initiating duplicate populations of each the following:

1. WT marked with Kan (40μl **from 1:1 WT Kan** into 4ml LB).
2. WT marked with Chl (40μl **from 1:1 WT Chl** into 4ml LB).
3. RpoB S512Y rifampicin-resistant marked with Kan (40μl **from 1:1 RpoB S512Y rifampicin-resistant Kan** into 4ml LB).
4. RpoB S512Y rifampicin-resistant marked with Chl (40μl **from 1:1 RpoB S512Y rifampicin-resistant Chl** into 4ml LB).
5. 10% WT marked with Kan combined with 90% RpoB S512Y rifampicin resistant marked with Chl (20μl **from 1:5 WT Kan** and 180μl **from 1:5 RpoB S512Y rifampicin-resistant Chl** into 3.8ml LB).
6. 10% WT marked with Chl combined with 90% RpoB S512Y rifampicin resistant marked with Kan (20μl **from 1:5 WT Chl** and 180μl **from 1:5 RpoB S512Y rifampicin-resistant Kan** into 3.8ml LB).
7. 1% WT marked with Kan combined with 99% RpoB S512Y rifampicin resistant marked with Chl (2μl **from 1:5 WT Kan** and 198μl **from 1:5 RpoB S512Y rifampicin-resistant Chl** into 3.8ml LB).
8. 1% WT marked with Chl combined with 99% RpoB S512Y rifampicin resistant marked with Kan (2μl **from 1:5 WT Chl** and 198μl **from 1:5 RpoB S512Y rifampicin-resistant Kan** into 3.8ml LB).

Each of the resulting 16 populations was then placed in a shaker incubator and allowed to grow at 225 rpm at 37°C, with daily one-hundred-fold (1:100, v/v) dilutions for 16 dilution cycles (days), for a total of a little over 100 generations of growth overall (1:100 dilution, allowing for ∼6.6 generations of growth. 6.6*16=105.6).

Each day, a subsample of each culture was mixed with glycerol in a final concentration of 50% (v/v) and archived at −80°C.

### Isolation of rifampicin-resistant bacteria from evolved mixed-resistance populations

Mixed-resistance populations contained both resistant and susceptible cells. To carry out competition experiments using evolved rifampicin-resistant cells, we first needed to isolate only resistant bacteria from these populations. To do so, the evolved populations that initially contained 90% or 99% rifampicin-resistant bacteria and that then underwent 16 dilution cycles were revived from their glycerol stocks and inoculated in 4 ml of fresh LB with rifampicin and either Kan or Chl (depending on the cassette with which resistant cells were marked within that population), in 14 ml tubes and allowed to grow for two hours at 225 rpm and 37°C to an OD_600_ of 0.2. The samples were diluted and plated in duplicate on LB agar plates supplemented with kanamycin (50 µg/ml), LB agar plates supplemented with chloramphenicol (25 µg/ml), and LB agar plates supplemented with rifampicin (100 µg/ml). Plates were incubated overnight, allowing for the growth of visible colonies, and colony-forming units (CFU) were then quantified by colony counting. This allowed us to verify that we were starting our competition with only resistant cells marked with the expected cassettes.

### Competition experiments

Competition experiments were carried out by growing two strains or population samples together, maintaining a close to 50:50 initial ratio, and then examining the ultimate ratio of each cell type following overnight growth. In each competition experiment, the two competing strains or population samples were grown separately to an OD of 0.4 and were then diluted 1:100 into the same 4 ml Luria Broth (LB) medium and incubated overnight at 37°C. To determine the initial relative frequencies of each strain, a sample from each initial population was diluted and plated in duplicate on LB agar plates, supplemented with either kanamycin (50 µg/ml) or chloramphenicol (25 µg/ml). Plates were incubated overnight, allowing for the growth of visible colonies, and colony-forming units (CFU) were then quantified by colony counting. If the initial relative frequencies of each strain were not approximately 50:50, the experiment was discarded. To minimize counting errors and noise in the data, whole experiments with plates containing over 400 colonies were discarded. Following the overnight incubation of the competing bacteria, the final relative abundances of each strain were determined as was done prior to the incubation, through plating on plates containing kanamycin or chloramphenicol and calculating CFU.

From the resulting initial and final CFU counts, the relative fitness was calculated according to:

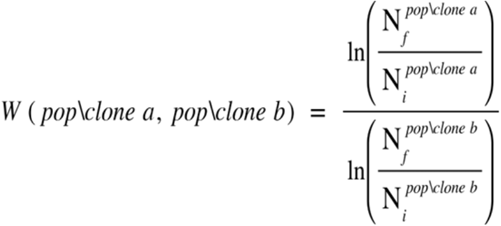

Where *W* is the relative fitness and *N* is CFU. The *i* and *f* subscripts denote the initial and final measurements, and the superscripts denote the population or clone of the WT and the mutated genes.

For each type of competition, at least ten individual experiments were carried out. The mean and the standard deviation around the mean were then calculated based on all individual experiments carried out for each type of competition, with these results being provided in **Tables 1** and **2**. N values and *W* values for each separate experiment are provided in **Tables S1** and **S2**.

Our measurements could be skewed if the cassettes exert very different effects on fitness. To make certain that this was not the case, both on the background of the susceptible (wildtype) ancestral strain and on the background of the ancestral rifampicin-resistant strain, we competed the Kan and Chl marked susceptible ancestral strains against each other and also competed the Kan and Chl marked resistant ancestral strains against each other. In both cases, we found no significant fitness effect of the cassettes themselves (**Table S5**).

### Sequencing and mutation calling

To extract clones for sequencing, frozen samples from each of the 12 sequenced populations that were archived at the end of the serial dilution evolutionary experiment (at day 16) were revived and grown overnight on LB agar plates. 10 colonies from each population were used to separately inoculate 4 ml of medium in a test tube and were grown until they reached an optical density of ∼1. This procedure was followed in order to minimize the number of generations each clone undergoes prior to sequencing and thus minimize the occurrence of mutations during regrowth. One milliliter of each clone’s culture was centrifuged at 5,000 g for 5 min, and the pellet was used for DNA extraction. The remainder of each culture was then archived by freezing in 50% of glycerol at −80 °C. DNA was extracted using the Macherey-Nagel NucleoSpin 96 Tissue Kit. Library preparation followed the protocol outlined in (29). Sequencing was carried out at Admera Health (New Jersey, USA) using an Illumina HiSeq machine. Clones were sequenced using paired-end 150 bp reads. The coverage obtained for each sequenced clone is provided in **Table S3**. Raw sequencing data were deposited in the Short Read Archive (SRA) under Bioproject ID PRJNA910842.

In order to call mutations, the reads obtained for each rifampicin-resistant clone were aligned to the *E*.*coli* K12 MG1655 reference genome (accession NC_000913). Mutations were then recorded if they did not appear within the susceptible ancestral genome, which was previously sequenced (Katz & Hershberg, 2013). Alignment and mutation calling were carried out using the Breseq platform, which allows for the identification of point mutations, short insertions and deletions, larger deletions, and the creation of new junctions (30). A full list of identified mutations is provided in **Table S4**.

### Simulations

We carried out simulations based on custom code in R (Supplementary text S1). The starting point is a population where the majority of cells carry a resistance mutation, and 1% or 10 % of the cells do not carry that mutation. The resistance mutation is associated with a fitness cost (c). We varied the fitness cost from 0.1 to 1 (in steps of 0.1) and varied the population size from 10^5^ to 10^10^. We set the mutation rate to 0. Next, we ran Wright-Fisher simulations (the population size is constant and multinomial sampling is used to determine the number of resistant and susceptible cells in the next generation). We then determined how many generations it takes for the resistant type to die out, given the cost of the mutation and the population size. In almost all cases, the resistant type will die out within 100 generations unless the cost of resistance is relatively low (0.1 or 0.2).

## Data availability statement

Raw sequencing reads were deposited to the sequence read archive (SRA) BioProject ID: PRJNA910842.

## Acknowledgments

This work was supported by an ISF grant No. 1860/21, to R.H and by the Rappaport Family Institute for Research in the Medical Sciences.

---

**Figure S1.**
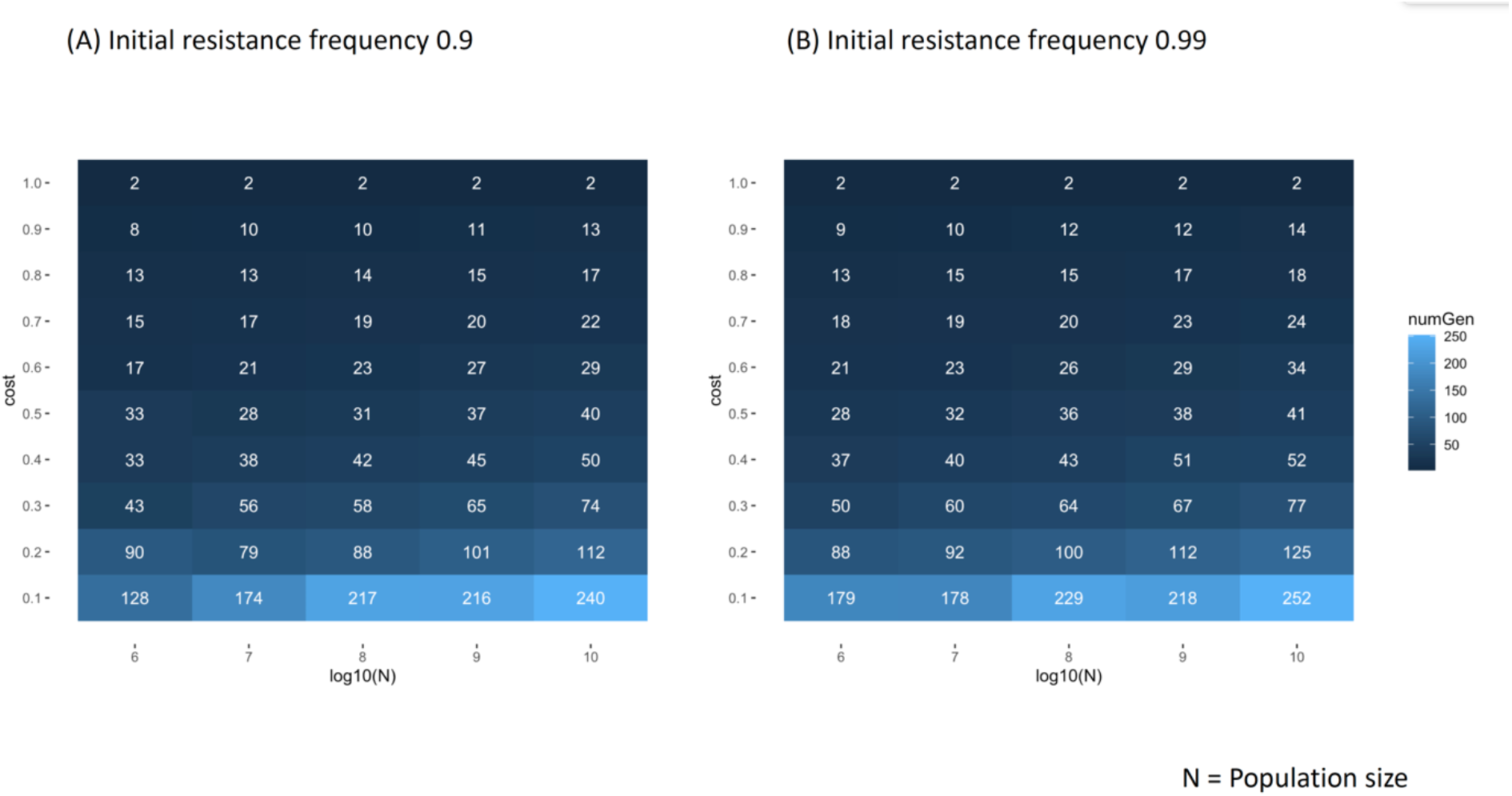
Simulation results for the quantification of the number of generations after which we would expect to observe a complete loss of resistant cells within populations evolving alongside susceptible cells, depending on the cost of resistance and population sizes. Two plots are given: (**A**) for an initial frequency of resistant cells of 0.9, and (**B**) for an initial frequency of resistant cells of 0.99.

